# Application of Passive Head Motion to Generate Defined Accelerations at the Heads of Rodents

**DOI:** 10.1101/2022.07.05.498781

**Authors:** Takahiro Maekawa, Naoyoshi Sakitani, Youngjae Ryu, Atsushi Takashima, Shuhei Murase, Julius Fink, Motoshi Nagao, Toru Ogata, Masahiro Shinohara, Yasuhiro Sawada

## Abstract

Exercise is widely recognized as effective for various diseases and physical disorders, including those related to brain dysfunction. However, molecular mechanisms behind the beneficial effects of exercise are poorly understood. Many physical workouts, particularly those classified as aerobic exercises such as jogging and walking, produce impulsive forces at the time of foot contact with the ground. Therefore, it was speculated that mechanical impact might be implicated in how exercise contributes to organismal homeostasis. For testing this hypothesis on the brain, a custom-designed passive head motion (hereafter referred to as PHM) system was developed that can generate vertical accelerations with controlled and defined magnitudes and modes and reproduce mechanical stimulation that might be applied to the heads of rodents during treadmill running at moderate velocities, a typical intervention to test the effects of exercise in animals. By using this system, it was demonstrated that PHM recapitulates the serotonin (5-hydroxytryptamine, hereafter referred to as 5-HT) receptor subtype 2A (5-HT_2A_) signaling in the prefrontal cortex (PFC) neurons of mice. This work provides detailed protocols for applying PHM and measuring its resultant mechanical accelerations at rodents’ heads.

**SUMMARY:** The present protocol describes a custom-designed passive head motion system, which reproduces mechanical accelerations at rodents’ heads generated during their treadmill running at moderate velocities. It allows dissecting mechanical factors/elements from the beneficial effects of physical exercise.

## INTRODUCTION

Exercise is beneficial to treat or prevent several physical disorders, including lifestyle diseases such as diabetes mellitus and essential hypertension^1^. Related to this, evidence has also been accumulated regarding the positive effects of exercise on brain functions^2^. However, molecular mechanisms underlying the benefits of exercise for the brain remain primarily unelucidated. Most physical activities and workouts generate mechanical accelerations at the head, at least to some extent. Whereas various physiological phenomena are mechanically regulated, the importance of mechanical loading has, in most cases, been documented in musculoskeletal system^3–5^. Although the brain is also subjected to mechanical forces during physical activities, particularly so-called impacting exercises, mechanical regulation of physiological brain function has rarely been studied. Because the generation of mechanical accelerations at the head is relatively common to physical workouts, it has been speculated that mechanical regulation might be implicated in the benefits of exercise to brain functions.

5-HT_2A_ receptor signaling is essential in regulating emotions and behaviors among various biochemical signals that function in the nervous system. It is involved in multiple psychiatric diseases^6–8^, on which exercise has been proven to be therapeutically effective. 5-HT_2A_ receptor is a subtype of 5-HT_2_ receptor that belongs to the serotonin family and is also a member of the G-protein-coupled receptor (GPCR) family, the signaling of which is modulated by its internalization, either ligand-dependent or -independent^9^. Head twitching is a characteristic behavior of rodents, the quantity (frequency) of which explicitly represents the intensity of 5-HT_2A_ receptor signaling in their prefrontal cortex (PFC) neurons^10,11^. Taking advantage of the strict specificity of this hallucinogenic response to administrated 5-HT (head-twitch response, hereafter referred to as HTR; see **Supplementary Movie 1**), the hypothesis mentioned above on mechanical implications in exercise effects on brain functions was tested. Thus, we analyzed and compared the HTR of mice subjected to either forced exercise (treadmill running) or exercise-mimicking mechanical intervention (PHM; **Supplementary Movie 2**).

## PROTOCOL

All animal experiments were approved by the Institutional Animal Care and Use Committee of National Rehabilitation Center for Persons with Disabilities. Eight- to nine-week old male Sprague-Dawley rats were used to measure accelerations at the head during treadmill running and PHM. Nine- to ten-week old male C57BL/6 mice were used for behavior tests and histological analyses of the PFC. The animals were obtained from commercial sources (see **Table of Materials**).

### 1. Measurement of magnitudes of accelerations along x-, y-, and z-axes during treadmill running

1.1. Anesthetize the rat with inhalation of 1.5% isoflurane. NOTE: The rats were used after at least 1 week of acclimation to the laboratory environment. Ensure that the rat is not responsive to a hindlimb toe pinch.
1.2. Fix the accelerometer (see **Table of Materials**) on top of the rat’s head using surgical tape.
1.3. After complete recovery from anesthesia, place the rat in the treadmill machine (see **Table of Materials**), and adjust the treadmilling to a moderate velocity (20 m/min)^12^ (**Figure 1A**). NOTE: It took at least 20 min to confirm the full recovery of the rat from anesthesia after the termination of isoflurane inhalation and start the treadmill experiment. Ensure that the rat is responsive to a hindlimb toe pinch, being able to walk or run without apparent staggering.
1.4. Measure the magnitude of vertical accelerations during rat treadmill running using the application software following the manufacturer’s instructions (see **Table of Materials**). NOTE: Extract 10 serial waves and individually calculate the average accelerations along the 3-dimensional axes (*x*-, *y*- *z*-axes, **Figure 1B**). Peak magnitudes were quantified by defining stepping-synchronized waves (^~^2 Hz frequency) as treadmill running-induced accelerations (**Figure 1C**).

**Figure 1:**
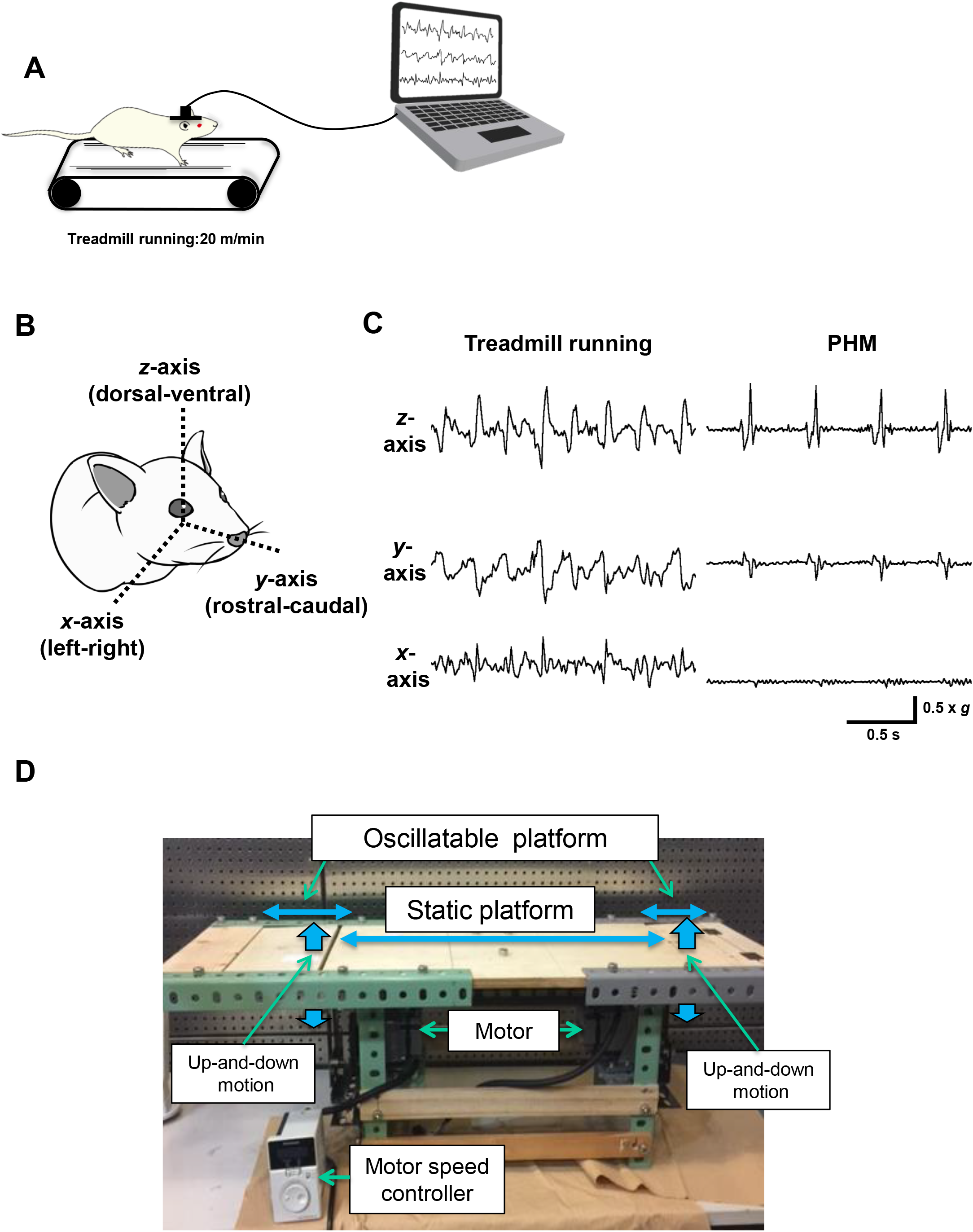
Measuring the magnitudes of accelerations during treadmill running. (**A**) Illustration for the measurement of accelerations generated at the heads of rats during their treadmill running. (**B**) Definition of x-(left-right), y-(rostral-caudal), and z-(dorsal-ventral) axes used in this study. (**C**) Accelerations were generated at rats’ heads during treadmill running at 20 m/min and PHM (frequency: 2 Hz) (n = 3 rats for each group). The PHM system was adjusted to produce vertical acceleration peaks equivalent to those during 20 m/min treadmill running (1.0 × *g*). Right-angled scale bar, 0.5 × *g* / 0.5 s. Images represent three independent experiments with similar results. (**D**) Photograph of the whole PHM system. (**E**) Photograph of propeller-shaped cam connected to a driver-equipped motor. (**F**) Photograph of propeller-shaped cam comprised of 4 blades with 5-mm step heights (see double-headed red arrow). The figure has been modified from Ryu et al.^15^.

### 2. Adjustment of the PHM system and application of PHM to mice

2.1. Pre-set the amplitude of the platform’s oscillation and the propeller-shaped cam’s rotation speed in the PHM system (**Figure 1D**) so the magnitude and frequency of vertical acceleration match the values obtained in step 1.4. NOTE: The PHM system is comprised of a metal framework and wooden platform. The motor speed can be changed and controlled by adjusting the dial connected to the build-in driver (see **Table of Materials**). The dial scale of 600 corresponds to 2 Hz, **Figure 1E**. The propeller-shaped cam has 4 blades with 5-mm step heights (**Figure 1F**).
2.2. Anesthetize a mouse *via* inhalation of 1.2% isoflurane. NOTE: Mice were used after at least 1 week of acclimation to the laboratory environments. Ensure that the mouse is not responsive to a hindlimb toe pinch.
2.3. Place a mouse in a prone position with the head and the rest of the body located on the oscillatable and static platforms, respectively. NOTE: Keep the mouse anesthetized (1.2% isoflurane).
2.4. Turn the motor on to oscillate the platform vertically, and apply PHM to the mouse. NOTE: The motor speed was adjusted to oscillate the platform at 2 Hz (see Protocol 2.1). Anesthetize and place the control mouse on the PHM platform likewise, but leave the motor off.

### 3. Running of a mouse on the treadmill

3.1. Place a mouse on the treadmill machine and adjust the treadmilling to a moderate velocity (10 m/min)^13^.

### 4. Quantification of mouse head-twitch response (HTR)

4.1. Set up a video camera (frame rate: 24 fps) to record the entire space in the transparent plastic case. Note: The plastic cage was used to keep the mouse in the field of video recording.
4.2. Intraperitoneally administer 5-hydroxytryptophan (5-HTP) (100 mg/kg) (see **Table of Materials**), the precursor to 5-HT, to a mouse.
4.3. Place the mouse in the transparent case and start recording for 30 min.
4.4. Review the recorded video (1/2× or 1/3 speed), counting the head twitching manually. NOTE: The analysts were not blinded to the experimental procedure. Characteristic “tic-like” rapid motion of mouse (see **Supplementary Movie 1**) was counted as head twitching, which rarely occurs under normal breeding environment.

### 5. Immunohistochemical analysis of mouse PFC

5.1. Once HTR tests are completed, anesthetize the mouse, perfuse with 4% paraformaldehyde (PFA) in PBS, and then excise the brain following previously published reports^14,15^.
5.2. Post-fix the brains in 4% PFA in PBS for an additional 24 h at 4 °C, and store in 30% sucrose/PBS until they sank. Freeze the fixed in optimal cutting temperature compound (OCT compound, see **Table of Materials**).
5.3. Retrieve the cryo-sections of the mouse brain from the slide box (see **Table of Materials**). Leave the slides on clean wipes (see **Table of Materials**) at room temperature until the samples dehydrate entirely. NOTE: Twenty-micrometer-thick sagittal sections (Lateral +0.5−1.5 mm) were prepared from frozen samples embedded in OCT compound using a cryostat (see **Table of Materials**).
5.4. Use a liquid blocker pen (see **Table of Materials**) to draw a circle around the cryo-sectioned tissue on the slide to confine the spread area of the solution (0.1% Tween-20 in Tris-buffered saline (TBS-T).
5.5. Place dampened wipes on the bottom of a tray holding the slides to create a moist environment.
5.6. After permeabilization with TBS-T, block with 4% donkey serum (see **Table of Materials**) at room temperature for 1 h.
5.7. Rinse the slides once by 5-min immersion in TBS-T.
5.8. Apply 100 μL of appropriately diluted primary antibody and DAPI (see **Table of Materials**) mix on each slide, cover the tray to avoid drying the sample, and incubate overnight at room temperature.
5.9. Rinse with TBS-T 3 times (5 min incubation each).
5.10. Apply 100 μL of appropriately diluted species-matched fluorescent secondary antibody (conjugated with Alexa Fluor 488, 568, or 645) (see **Table of Materials**) on each slide and incubate for 1 h at room temperature.
5.11. Rinse with TBS-T 3 times (5 min incubation each).
5.12. Mount the slides with mounting medium (see **Table of Materials**). Cover the slides with coverslips.
5.13. View the sample with a fluorescence microscope.

## REPRESENTATIVE RESULTS

The peak magnitude of the vertical accelerations at rats’ heads during their treadmill running at a moderate velocity (20 m/min) was approximately 1.0 × *g* (**Figure 1C**). The PHM system (**Figure 1D**) was set up to generate vertical acceleration peaks of 1.0 × *g* at rodents’ heads.

PHM application (2 Hz, 30 min/day for seven days) to mice significantly attenuated their HTR as compared to the control mice (daily anesthetized without PHM for 30 min/day for seven days) (**Figure 2**). This represents a suppressive effect of PHM on 5-HT_2A_ receptor signaling in the PFC neurons.

**Figure 2:**
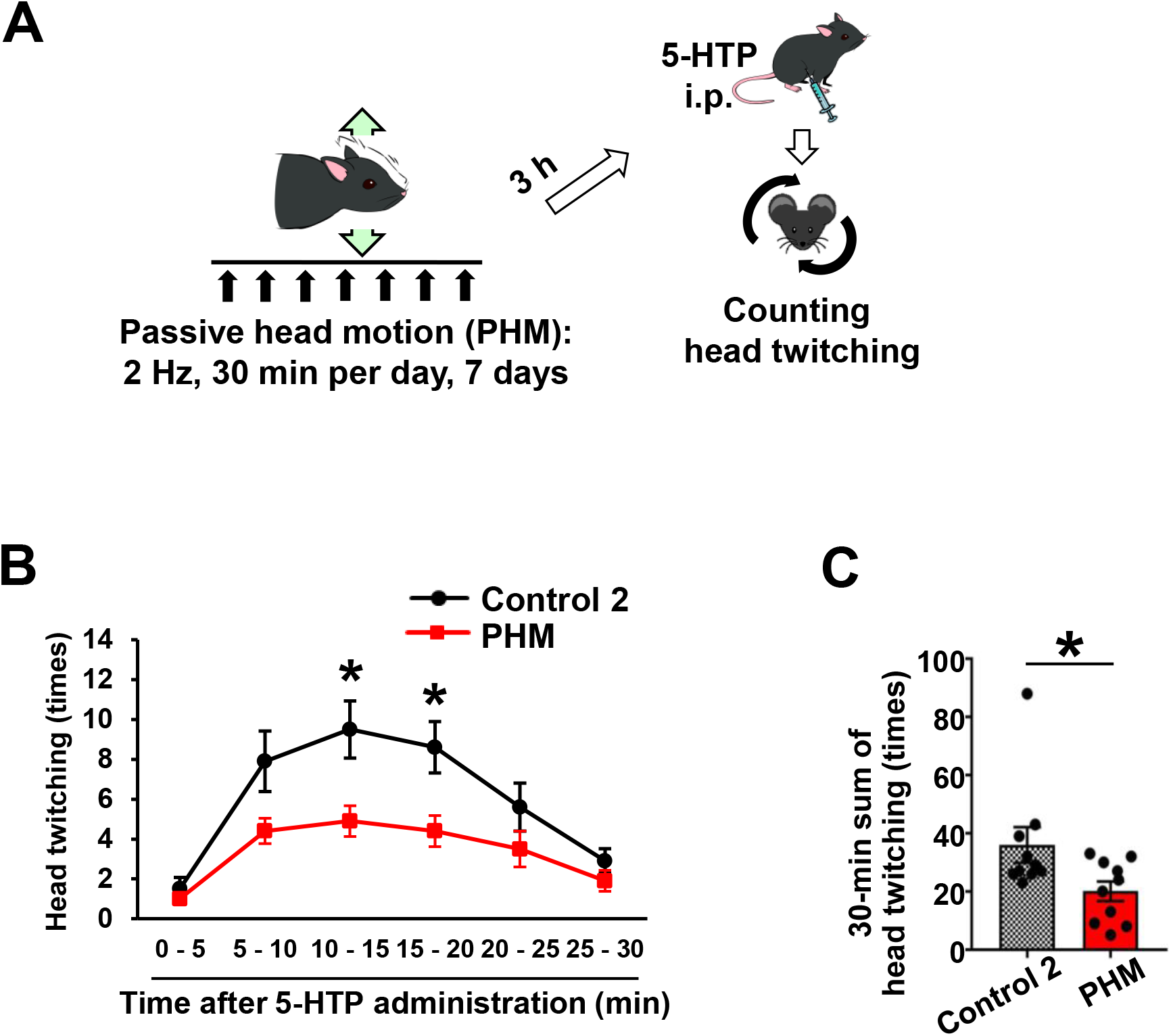
PHM application to mice. (**A**) Illustration for the analysis of the effects of PHM on HTR. (**B,C**) PHM mitigated 5-HTP-induced HTR. Head twitching was counted in 5-min blocks (**B**) and 30 min (**C**) after 5-HTP administration. Control 2 represents mice that were anesthetized and placed on the PHM platform left unoscillated. Data are presented as means ± SEM. *, *P*<0.05, unpaired t-test (n = 10 mice for each group). The figure has been modified from Ryu et al.^15^.

The treadmill running and PHM significantly enhanced 5-HT_2A_ receptor internalization in mouse PFC neurons (**Figure 3**). Consistently, both treadmill running and PHM down-regulated 5-HTP-induced c-Fos expression, the downstream cellular event of 5-HT_2A_ receptor activation^14^, in mouse PFC neurons (**Figure 4**). These results suggest that treadmill running and PHM internalize 5-HT_2A_ receptors in the PFC neurons, attenuating relevant signaling.

**Figure 3:**
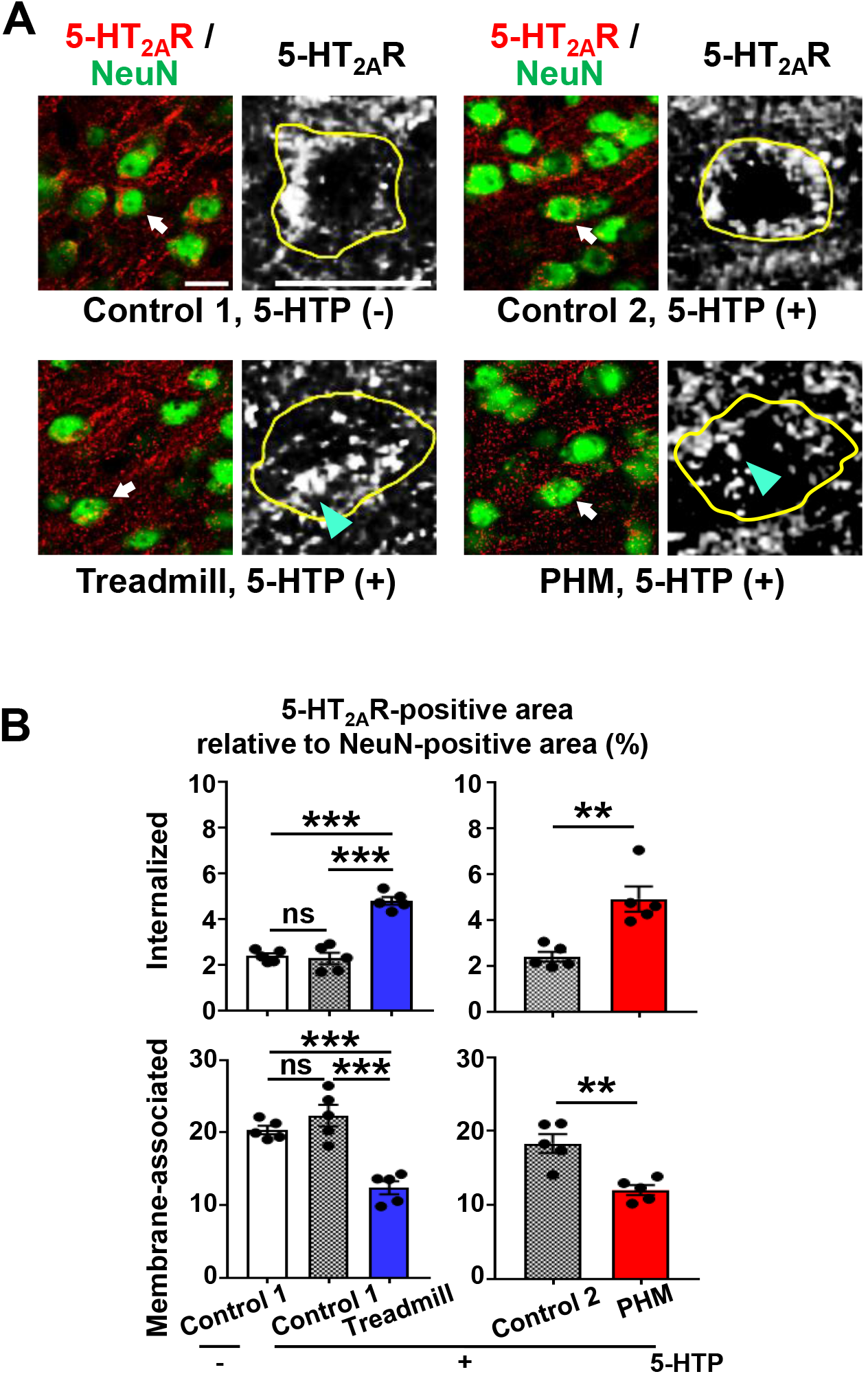
The treadmill running and PHM application enhanced 5-HT_2A_ receptor internalization in mouse PFC neurons. (**A**) Micrographs of anti-5-HT2A receptor (5-HT2_A_R; red) and anti-NeuN (green) immunostaining of the PFC of mice injected with 5-HTP (or vehicle) after a week of daily PHM. Higher-magnification images of anti-5-HT_2A_ receptor immunostaining of arrow-pointed cells are shown with a grayscale. Yellow lines indicate the soma margins outlined by NeuN-positive signals, and cyan arrowheads point to internalized anti-5-HT_2A_ receptor immunosignals. Scale bars, 20 μm. Images are representative of five mice. (**B**) Quantification of 5-HT_2A_ receptor internalization in mouse PFC neurons. Internalized and membrane-associated 5-HT_2A_ receptor-positive areas were quantified as values relative to the NeuN-positive area in mouse PFC. Control 1 represents mice placed in the treadmill machine left turned off, and control 2 represents mice that were anesthetized and placed on the PHM platform left unoscillated. Thirty-five to forty NeuN-positive neuronal somas were analyzed for each mouse (Internalized: left chart, p<0.001, one-way ANOVA with post hoc Bonferroni test; right chart, *P* = 0.0027, unpaired t-test; Membrane-associated: left chart, *P*<0.001, one-way ANOVA with post hoc Bonferroni test; right chart, *P* = 0.0025, unpaired t-test; n = 5 mice for each group). Data are represented as means ±SEM. \*\**P*<0.01, \*\*\**P*<0.001; ns, not significant. The figure has been modified from Ryu et al.^15^.

**Figure 4.**
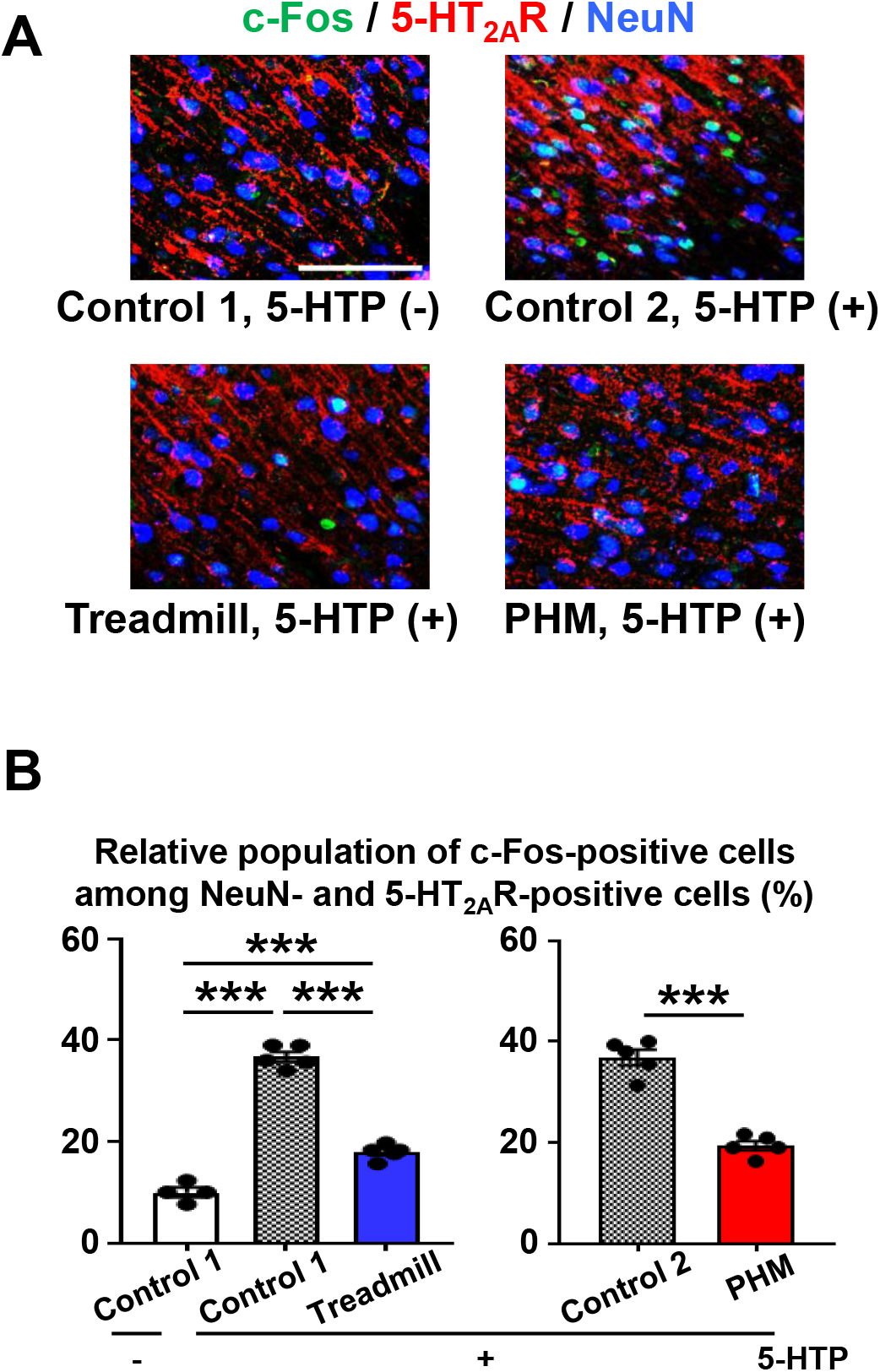
The treadmill running and PHM application down-regulated 5-HTP-induced c-Fos expression in mouse PFC neurons. (**A**) Micrographs of anti-c-Fos (green), anti-5-HT2A receptor (red), and anti-NeuN (blue) immunostaining of the PFC of mice intraperitoneally administered with 5-HTP (or vehicle) after a week of daily PHM. Scale bar, 100 μm. Images are representative of four to five mice. (**B**) Quantification of c-Fos expression in 5-HT_2A_ receptor-positive neurons in mouse PFC. Control 1 represents mice placed in the treadmill machine left turned off, and control 2 represents mice that were anesthetized and placed on the PHM platform left unoscillated. Relative population (%) of c-Fos-positive cells of 300 NeuN- and 5-HT_2A_ receptor-positive cells is shown (left chart: *P*<0.001, one-way ANOVA with post hoc Bonferroni test; right chart: *P*<0.001, unpaired t-test; n = 4 mice for columns 1, n = 5 mice for columns 2 to 5). Data are represented as means ±SEM. \*\*\**P*<0.001. The figure has been modified from Ryu et al.^15^.

---

**Supplementary Movie 1: Head-twitch response of a mouse.** The 2-min 46-sec movie starts 6 min after the HTP injection. Head twitching is observed at the time points of 0:03, 0:39, 1:39, and 2:42.

**Supplememtary Movie 2: Mouse PHM.** (**A**) Side view. (**B**) Top view.

## DISCUSSION

Using the developed PHM application system, we have shown that 5-HT signaling in their PFC neurons is mechanically regulated. Because of the complexity of exercise effects, it has been difficult to precisely dissect the consequence of exercise in the context of health promotion. The focus is on mechanical aspects to preclude the involvement or contribution of metabolic events that may occur with or subsequently to exercise activities, such as energy consumption. The method described here is expected to be more broadly useful in biomedical research exploring the mechanisms underlying exercise effects on brain functions.

The current system requires anesthesia to subject the experimental animals to PHM, which may influence (either detrimental or not) nervous cell behavior and processes in the brain. It may be possible to apply PHM without anesthesia by modifying the system, including the magnitudes, modes, and wave shapes of mechanical accelerations generated by PHM. For example, smaller accelerations peaks with sinusoidal waves instead of 1 × *g* impulsive peaks of the current PHM may be “felt” as more comfortable stimulation by the animals. Alternatively, new method(s) can be implemented to hold experimental animals on the oscillating platform with minimal stress. These modifications and improvements are feasible, mainly because the PHM-generated accelerations relate to moderate exercise in principle and are unlikely to be “painful” stress for experimental animals.

Many previous studies have reported moderate exercise as an effective procedure to treat or prevent numerous diseases and disorders^16,17^. Lactate threshold, where plasma lactate concentration increases exponentially with adrenocorticotrophic hormone (ACTH) secretion, a stress indicator^18^, is utilized to determine exercises as mild or moderate^19^. However, “optimal” exercise remains to be defined at a molecular level. Because not only the brain but eventually all the other bodily organs are subjected to mechanical forces during exercise, the current approach employing mechanical perturbations may be helpful to reveal the molecular mechanisms behind the exercise effects in broader contexts and help define “what is optimal exercise” by scientific measures.

However, the current method suffers certain limitations. We could not stably fix the accelerometer on the mouse head because of the lack of size compatibility. Although the preliminary measurement indicates that the peak magnitude of treadmill running-generated mechanical accelerations at the mouse head is also approximately 1.0 × *g*, further studies are required to quantify it more accurately.

The present protocol has detailed the procedures of the custom-designed PHM system, which allowed for dissecting mechanical elements/factors from physical exercise. The approach provides significant insights into the benefits of exercise to brain functions.

## ACKNOWLEDGMENTS

This work was in part supported by the Intramural Research Fund from the Japanese Ministry of Health, Labour and Welfare; Grants-in-Aid for Scientific Research from the Japan Society for the Promotion of Science (KAKENHI 15H01820, 15H04966, 18H04088, 20K21778, 21H04866, 21K11330, 20K19367; MEXT-Supported Program for the Strategic Research Foundation at Private Universities, 2015-2019 from the Japanese Ministry of Education, Culture, Sports, Science and Technology (S1511017); the Naito Science & Engineering Foundation.

