## Supplementary material for "Application of Passive Head Motion to Generate Defined Accelerations at the Heads of Rodents": Table of Materials

| Name of Material/Equipment | Company | Catalog Number | Comments/Description |
|---|---|---|---|
| 5-hydroxytryptophan (5-HTP) | Sigma-Aldrich | H9772 | Serotonin (5-HT) precursor |
| Brushless motor driver | Oriental motor | BMUD30-A2 | Speed changer build-in motor driver |
| C57BL/6 mice | Oriental yeast company | C57BL/6J | Mice used in this study |
| Cryostat | Leica | CM33050S | Microtome to cut frozen samples |
| DC Motor | Oriental motor | BLM230-GFV2 | Motor |
| Donkey anti-goat Alexa Fluor 568 | Invitrogen | A-11057 | Secondary antibody used for immunohistochemical staining |
| Donkey anti-mouse Alexa Fluor 647 | Invitrogen | A-31571 | Secondary antibody used for immunohistochemical staining |
| Donkey anti-rabbit Alexa Fluor 488 | Invitrogen | A-21206 | Secondary antibody used for immunohistochemical staining |
| Donkey serum | Sigma-Aldrich | S30-100ML | Blocker of non-specific binding of antibodies in immunohistochemical staining |
| Fluorescence microscope | Keyence | BZ-9000 | Fluorescence microscope |
| Goat polyclonal anti-5-$HT_{2A}$ receptor | Santa Cruz Biotechnology | sc-15073 | Primary antibody used for immunohistochemical staining |
| Isoflurane | Pfizer | v002139 | Inhalation anesthetic |
| KimWipe | NIPPON PAPER CRECIA | S-200 | Paper cloth for cleaning surfaces, parts, instruments in labratory |
| Liquid Blocker | Daido Sangyo | PAP-S | Marker used to make the slide surface water-repellent |
| Mouse monoclonal anti-NeuN (clone A60) | EMD Millipore (Merck) | MAB377 | Primary antibody used for immunohistochemical staining |
| NinjaScan-Light | Switchscience | SSCI-023641 | Accelerometer to measure accelerations |
| OCT compound | Sakura Finetek | 45833 | Embedding agent for preparing frozen tissue sections |
| ProLong Gold Antifade Mountant | Invitrogen | P36934 | Mounting medium to prevent flourscence fading |
| Rabbit polyclonal anti-c-Fos | Santa Cruz Biotechnology | sc-52 | Primary antibody used for immunohistochemical staining |
| Slide box | AS ONE | 03-448-1 | Opaque box to store slides |
| Spike2 | Cambridge electronic design limited (CED) | N/A | Application software used to analyze acceleration |
| Sprague-Dawley rats | Japan SLC | Slc:SD | Rats used in this study |
| Treadmill machine | Muromachi | MK-680 | System used in experiments of forced running of rats and mice |
